# “*In silico* characterization of Protein L-Iso-Aspartate-O-Methyltransferases (PIMT) of *Shigella flexneri*”

**DOI:** 10.1101/2023.07.03.547495

**Authors:** Akanksha Gupta, Pragati Mardi, Saumya, Prasanta Kumar K. Mishra

## Abstract

There are only a few protein repair enzymes known. One of them, Protein Iso-Aspartate Methyltransferases (PIMT), also known as Protein Carboxyl Methyltransferases (PCMT), is responsible for converting Iso-Aspartate (abnormal amino acid residues) to Aspartate (normal amino acid residues). PCMT is an enzyme found in a wide range of living organisms. In the present study, the amino acid sequence of PIMT of *Shigella flexneri* was retrieved and the 3D structure is predicted. Further, it is characterized by *in silico* approaches and its interaction with one of the target proteins *i. e*. rpoS is studied.

## Introduction

Misfolded, unfolded, and abnormal proteins are formed as a result of errors in the post translational modification of proteins and the malfunction of chaperone machinery. Proteins are constantly damaged by a variety of extrinsic and intrinsic factors. Sugars, aldehydes, reactive nitrogen and oxygen species, as well as oxidative modification of their lateral chain of amino acids, can harm them. Many proteins’ conformational stability and activity are affected as a result of this. To prevent the accumulation of abnormal proteins, the majority of damaged proteins are degraded by the proteasome system or repaired by protein repair enzymes like PIMT. Methylation of proteins is a complex and multifaceted process **[1]**. Protein Methyltransferases (PMTs) are the enzymes that catalyse the methylation of proteins **[2]**. S-adenosylmethionine (SAM) is used as a methyl group donor by protein methyltransferase enzymes **[3,4]**. Polypeptides are sensitive to a range of spontaneous, structural modifications that can cause structural and biological activity to be impacted. By regulating histone tail modification, PMTs play a functional role in cell response in stressful situations and protein repair **[5]**. Deamidation and dehydration of Asn and Asp of amino acid are examples of protein damage. Under oxidative stress, the conversion of Asparginyl (Asn) and Aspartyl (Asp) residues to L-Isoaspartyl (IsoAsp) residues as a major compound occurs via the formation of the L-Succinimide intermediate **[6,7]**. The nucleophilic attack of the adjacent amino acid nitrogen on the side chain carbonyl stimulates the alteration of Asp and Asp residues, resulting in the formation of a cyclic ring between the side chain and the main chain, resulting in the formation of a succinimide intermediate **[8]**.

## Material methods

### Sequence retrieval and characterization of the protein

The amino acid sequence of the protein PIMT was retrieved from NCBI database (WP_000254711.1). The target species was selected as *S. flexneri*. The amino acid sequence was analysed by ProtopParam tool of Expasy server (https://web.expasy.org/protparam/)and the result is presented in a tabular form. The amino acid sequence was further searched for presence of any transmembrane domain by TMHMM prediction (https://services.healthtech.dtu.dk/services/TMHMM-2.0/). The interactome of the protein was predicted by string server (https://string-db.org/cgi/network).

### Prediction of 3D structure of the protein

The amino acid sequence of the protein was submitted to PHYRE 2.0 server. Default parameters were opted and the search was performed in ‘normal mode”. The predicted structure was retrieved as a .pdb file and visualized by chimera software [9]. The single residues are named and shown in the structure. The 2D structural components were marked by different colours. The Ramachandran’s plot was predicted by using an online server.

### Determination of the active site of the enzymes

The amino acid sequence was submitted in .pdb format to the http://www.scfbio-iitd.res.in/server. The binding cavities were retrieved and presented in a tabular form.

### Docking studies

The 3D structure of the rpoS protein (5H6X) was taken from RCSB database (https://www.rcsb.org/). HEX 8.0 was used for docking. The frame rate was fixed to 35 and rest default parameters were opted for the analysis.

## Results and discussion

The PIMT amino acid sequence retrieved was of 208 residues. The theoretical pI of the protein was found to be 6.53. The average molecular weight of the protein was found to be 23.2 kDa. The instability index of the protein was found to be 49 which is an indication that the protein is unstable unlike many of its homologs present in other species. The characteristic details are given as table. 1.

**Table 1:** The biochemical properties of PIMT of *Shigella flexneri*

| Properties | Value |
| --- | --- |
| Number of amino acids | 208 |
| Molecular weight | 23.2 kDa |
| Theoretical pI | 6.5 |
| Total number of atoms | 3303 |
| The estimated half-life is | 30 hours (mammalian reticulocytes, in vitro). >20 hours (yeast, in vivo). >10 hours (Escherichia coli, in vivo). |
| Aliphatic index | 101.73 |
| Grand average of hydropathicity (GRAVY) | -0.183 |

PIMT has been found in the cytoplasmic fraction in the species reported till date. However, in bacteria, the reports are few [10]. In *S. flexneri*, it was found that there is no transmembrane domain present (Fig. 1). But interestingly, the residues were mostly found to remain outside. It may indicate the possibility of such proteins to be secretory.

**Fig. 1:**
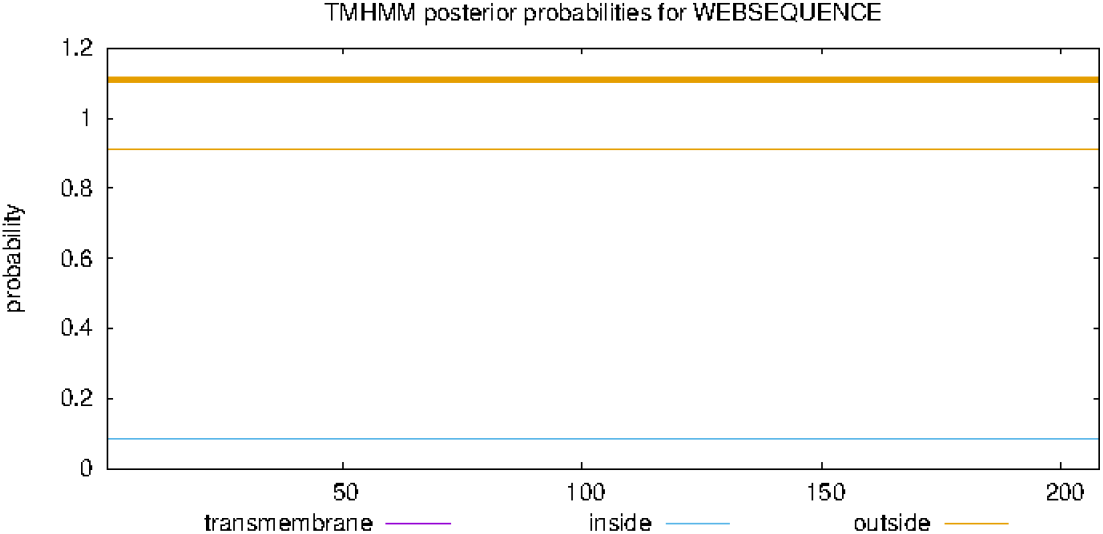
The predicted residues of PIMT of the ***S. flexneri*** for spanning the membrane or distribution across the membrane

Further, analysing the interactome of the PIMT suggested that it interacts with cmr, gltB, ygbB, Lipoprotein, surE, rpoS etc (Fig. 2).

**Fig. 2:**
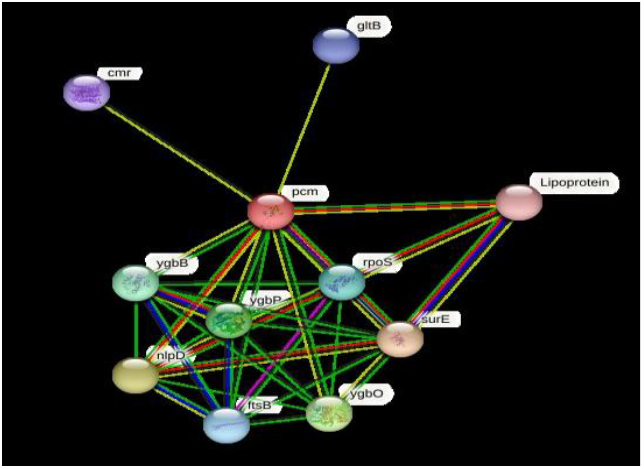
The interactome of PIMT of the ***S. flexneri*** as predicted by String server.

The predicted 3D structure is presented in fig. 3. The one letter codes of amino acids are given to their respective positions. The secondary structure i. e. α-helix and β-sheets are marked in the fig. 4. The Ramachandran’s plot is given in the fig. 5. This indicates an equal presence of both helix and sheets in the protein structure of the *Shigella* PIMT.

**Fig. 3:**
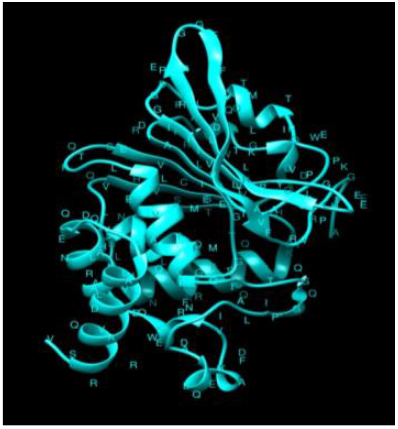
The 3D structure of PIMT of the ***S. flexneri***. The structure was visualized by Chimera software.

**Fig. 4:**
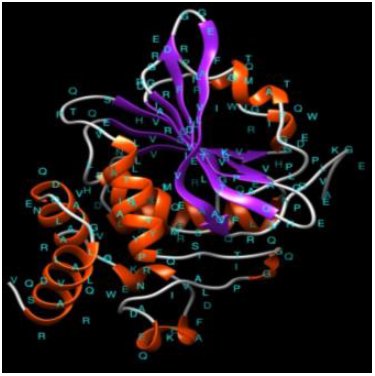
The 2° structural content of PIMT of the ***S. flexneri***. The Red coloured area represents helixes while purple colour represents antiparallel sheets.

**Fig. 5:**
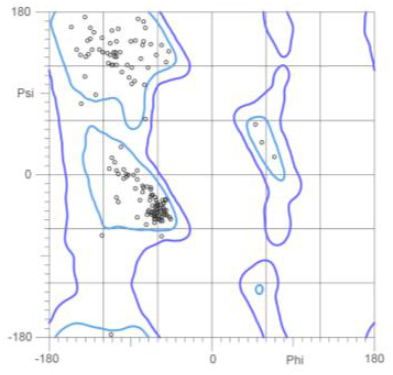
The Ramachandran plot of PIMT of the ***S. flexneri***.

The docking sites of the enzyme were predicted as in the form of cavities. The sequence of the 12 potential cavities are given in table 2.

**Table 2:**
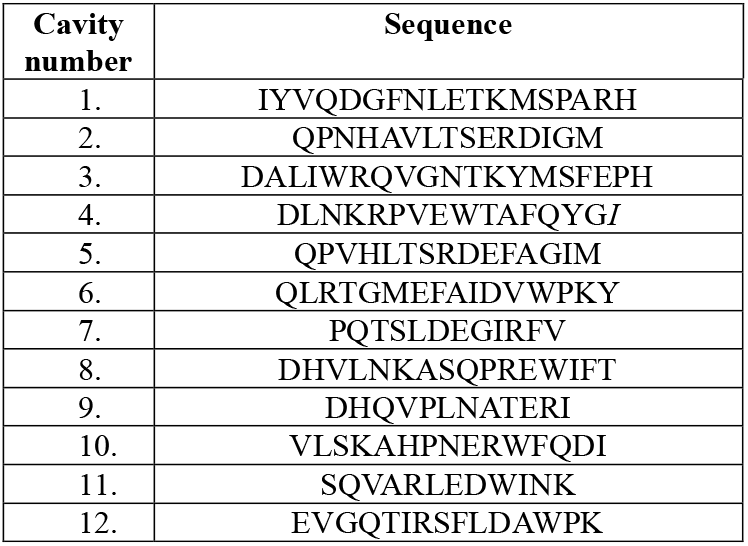
Binding cavities of PIMT

Both the proteins were docked taking PIMT as the receptor and rpoS as ligand. The E total was found to be -416.46, E – shape as -541.26, E-force as-124.80. An image of the docked complex is presented as fig 6.

**Fig. 6:**
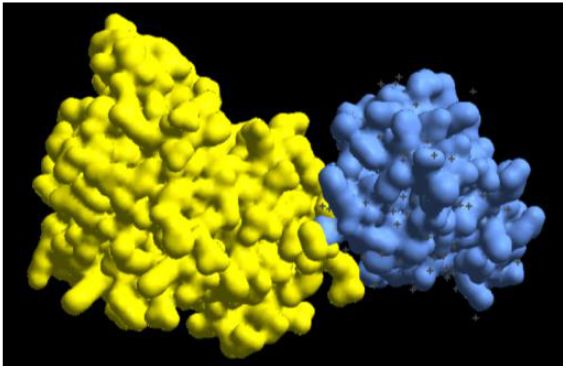
PIMT docked with rpoS

## Conclusions

The PIMT is an interesting target for vaccine candidate, drugs targeting ageing, plant survival and seed vigour etc. Various efforts must be made to elucidate its function and new targets. The present study though has described about the in-silico characteristics of the enzyme but invitro analysis are needed to corroborate the findings.

